# A deep mutational scan of an acidic activation domain

**DOI:** 10.1101/230987

**Authors:** Max V. Staller, Alex S. Holehouse, Devjanee Swain-Lenz, Rahul K. Das, Rohit V. Pappu, Barak A. Cohen

## Abstract

Transcriptional activation domains are intrinsically disordered peptides with little primary sequence conservation. These properties have made it difficult to identify the sequence features that define activation domains. For example, although acidic activation domains were discovered 30 years ago, we still do not know what role, if any, acidic residues play in these peptides. To address this question we designed a rational mutagenesis scheme to independently test four sequence features theorized to control the strength of activation domains: acidity (negative charge), hydrophobicity, intrinsic disorder, and short linear motifs. To test enough mutants to deconvolve these four features we developed a method to quantify the activities of thousands of activation domain variants in parallel. Our results with Gcn4, a classic acidic activation domain, suggest that acidic residues in particular regions keep two hydrophobic motifs exposed to solvent. We also found that the specific activity of the Gcn4 activation domain increases during amino acid starvation. Our results suggest that Gcn4 may have evolved to have low activity but high inducibility. Our results also demonstrate that high-throughput rational mutation scans will be powerful tools for unraveling the properties that control how intrinsically disordered proteins function.

## Introduction

Transcription factors (TFs) activate target genes by binding to specific DNA sequences with DNA binding domains (DBDs) and recruiting coactivator complexes with activation domains (Latchman, 2008). While DBDs tend to be phylogenetically conserved, well structured, and bind to related DNA sequences, activation domains are poorly conserved, intrinsically disordered and bind structurally diverse coactivators (Dyson and Wright, 2016; Hahn and Young, 2011; Latchman, 2008). When a new genome is sequenced, homology-based models for known DBDs can identify candidate TFs (Finn et al., 2011, 2016). However, because we lack robust models of activation domains, we cannot predict whether new TFs are activators, repressors, or both. Moreover, it is hard to identify activation domains on regulatory proteins without traditional DNA binding domains (e.g. Notch). Even in known activation domains, we cannot predict the functional consequences of mutations. As a result, the majority of clinically identified variants in oncogenic and tumor suppressing TF activation domains are classified as variants of uncertain significance (Forbes et al., 2015). The gap in understanding between DBDs and activation domains remains a critical challenge in the study of gene regulation.

Our understanding of DBDs has benefited from high-throughput methods for measuring protein-DNA interactions *in vitro* (Bulyk, 2007; Riley et al., 2014; Stormo et al., 2015; Teytelman et al., 2013) and *in vivo* (Buenrostro et al., 2013; Noyes et al., 2008; Vierstra and Stamatoyannopoulos, 2016). The resulting data provide statistical power to populate sophisticated computational models of DNA-protein interactions (Riley et al., 2015; Stormo and Zhao, 2010) and in some cases predict DNA-affinity directly from an amino acid sequence (Persikov et al., 2015). No analogous high-throughput methods exist for studying activation domains. Consequently, we lack computational models for predicting activation domains from amino acid sequences.

The current working models of activation domains are supported by conflicting data. The first few eukaryotic examples were all negatively charged, leading to the ‘acid blob’ model, wherein an activation domain is a cluster of negatively charged residues (Gill and Ptashne,1987; Hope et al., 1988; Ma and Ptashne, 1987a, 1987b; Sigler, 1988). However, directed mutagenesis showed that acidic residues often have only a small effect on activity and instead the most important positions are bulky hydrophobic residues (Cress and Triezenberg, 1991; Drysdale et al., 1995; Regier et al., 1993). With rare exceptions (Chen et al., 2008), all structurally characterized activation domains are intrinsically disordered regions (IDRs) and some form helices when bound to coactivators (Brzovic et al., 2011; Dyson and Wright, 2016; Jonker et al., 2005; Uesugi et al., 1997). Preformation of a helix can increase affinity for a binding partner (Borcherds et al., 2014; Warfield et al., 2014) and predicted helical propensity can correlate with activity (Scholes and Weinzierl, 2016). However, the necessity of helix formation for function has been questioned because adding a helix-breaking proline typically causes only a modest reduction in activity (Brzovic et al., 2011; Cress and Triezenberg, 1991). One unified model for all activation domains is that they are comprised of short linear motifs of hydrophobic residues embedded in IDRs (Warfield et al., 2014). These motifs are frequently presented to binding partners on one side of an α-helix (Dyson and Wright, 2016), reminiscent of the ‘amphipathic helix’ model (Giniger and Ptashne, 1987).

The sequence features that control activation domain function have been hard to resolve because the implicated properties are related. Intrinsic disorder and hydrophobicity are anticorrelated in the proteome and highly acidic regions tend to be intrinsically disordered. Since most mutations simultaneously change acidity, hydrophobicity, and intrinsic disorder, the relative contributions of these properties and short peptide motifs remain unknown. Resolving this problem requires experiments specifically designed to disentangle the effects of these convoluted variables.

In this work, we chose as our model a classic acidic activation domain, the central region of the yeast TF, Gcn4 (residues 101-144). Although this region was one of the first acidic activation domains identified, mutagenesis showed that the acidic residues are not essential for activity, but that the hydrophobic residues are (Drysdale et al., 1995; Hope and Struhl, 1986; Hope et al., 1988). The Gcn4 activation domain is intrinsically disordered, but folds into a helix when it binds the coactivator, Gal11 (Brzovic et al., 2011). However, introducing a helix-breaking proline leads to only a modest decrease in activity (Brzovic et al., 2011). At the center of this helix lies a well characterized hydrophobic motif, WxxLF (Brzovic et al., 2011; Drysdale et al., 1995; Warfield et al., 2014), which contributes to its activity.

To examine the sequence features that control the function of this activation domain, we developed a high-throughput fluorescent reporter assay to measure the strength of thousands of rationally designed mutants in the Gcn4 activation domain. Each variant contains 4-6 substitutions that independently alter either the charge, hydrophobicity or predicted disorder of the Gcn4 activation domain. We demonstrate that the activity of the Gcn4 activation domain increases during amino acid starvation, revealing a new layer of induction on top of known translational induction during starvation (Hinnebusch, 2005). Most mutations increased baseline activity and decreased induction, suggesting that Gcn4 may have been selected for low activity and high inducibility. We show how activity and induction are controlled by the number and placement of key aromatic residues in a disordered peptide scaffold with appropriately positioned acidic residues. Taken together our data support a unified model of activation domain function in which acidic residues and conformational disorder keep key hydrophobic residues exposed to solvent and binding partners. We conclude that deep, but rational mutational schemes will be an important set of tools for unraveling the sequence features that define transcriptional activation domains.

## Results

### A high-throughput reporter assay for quantifying activation domain strength

We present a high-throughput reporter assay for measuring the activities of thousands of rationally designed mutations in transcriptional activations domains (Figure 1A). The library of designed activation domain mutants is expressed in budding yeast as part of a synthetic TF. The TF construct contains 1) mCherry (to normalize for the abundance of each variant (Shaner et al., 2004)), 2) the murine Zif268 DBD (to avoid interfering with native yeast gene regulation (McIsaac et al., 2013a)), 3) a human estrogen response domain (to make the library inducible with ß-estradiol (McIsaac et al., 2013a)), 4) a rationally designed Gcn4 activation domain mutant (described below) and 5) a unique barcode sequence in the 3’ UTR that marks the identity of each library member (Methods). The library is integrated into the genome at a single locus to reduce expression variability and to ensure one variant per cell.

**Figure 1:**
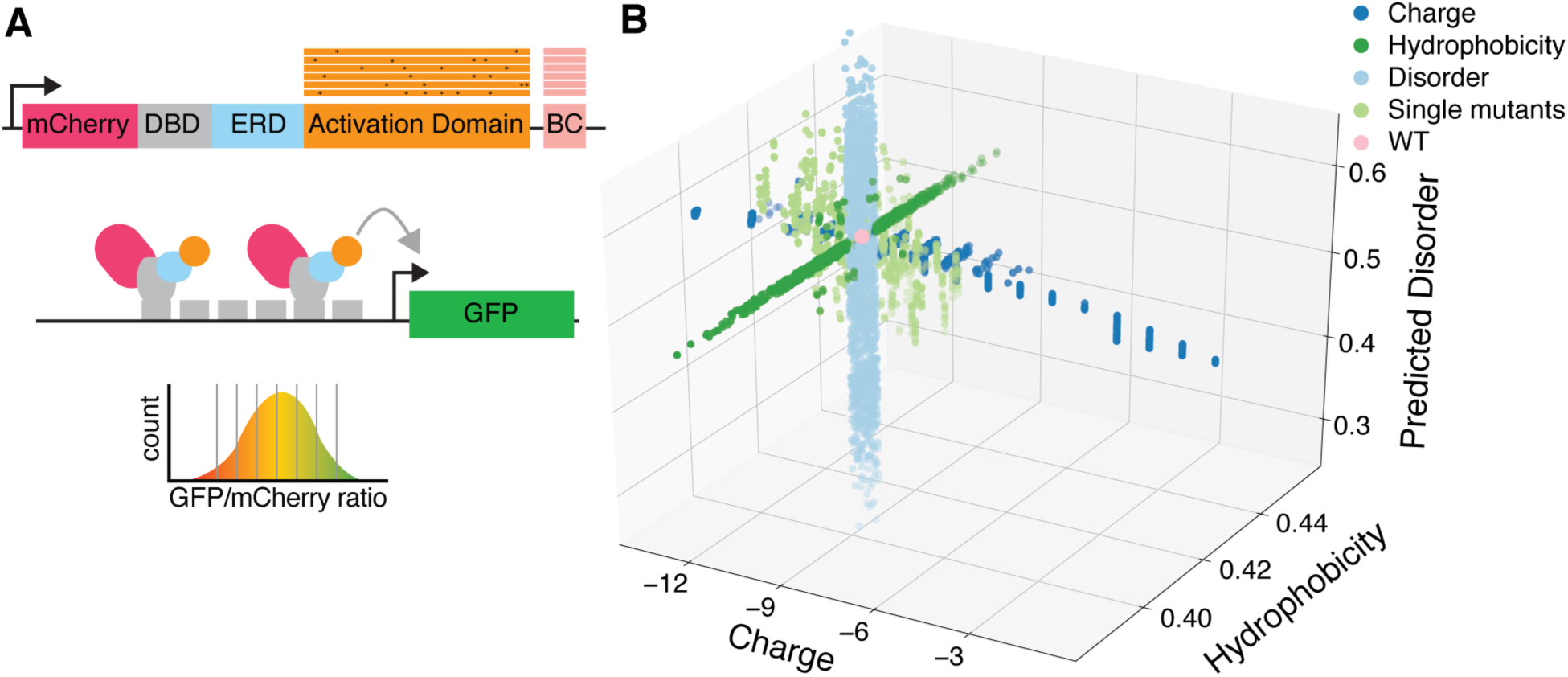
A method for measuring thousands of designed activation domain mutants in parallel. (**A**) The method for assaying activation domain variants uses a synthetic TF and a reporter. The synthetic TF contains mCherry, the murine Zif268 DNA-binding Domain (DBD), a human estrogen response domain (ERD), a designed activation domain variant and a barcode (BC). Each cell contains one TF variant integrated into the chromosome. The TF binds the promoter of a GFP reporter containing six binding sites for the DBD. Functional activation domains will activate the GFP reporter. Cells are sorted based on the GFP to mCherry ratio, genomic DNA is extracted from each bin and barcodes are sequenced. The relative abundance of each barcode across the bins is multiplied by the median ratio of each bin to calculate an activity. (**B**) Each point is a variant of the Gcn4 activation domain plotted according to its net charge, net hydrophobicity and predicted disorder. Mutations were rationally designed to change one physical property at a time. e.g. the disorder mutants (light blue) do not change charge or hydrophobicity. These designed mutations span a larger range of physical parameters than a mutational scan (light green) where each position is mutated to all 19 other amino acids.

The activity of each library member is measured using a genome-integrated reporter gene containing six copies of the Zif268 binding site driving GFP (McIsaac et al., 2013a). In each experiment, we induce nuclear localization of the TF library with ß-estradiol and use Fluorescence Activated Cell Sorting (FACS) to sort cells into bins according to the ratio of GFP to mCherry fluorescence. Highly active mutants have a high GFP to mCherry ratio, while inactive variants have a low ratio. We quantify the abundance of each variant in each bin by harvesting genomic DNA from each bin and sequencing barcodes. Using the relative abundance of each barcode across the bins and the median fluorescence ratio of each bin, we compute an *activity* for each variant (Methods) (Kinney et al., 2010; Sharon et al., 2012). Biological replicates are highly reproducible (Pearson correlation (*R_P_*) > 0.98, Spearman correlation (*R_s_*) > 0.98) and neither abundance in the library, nor protein abundance bias activity (**Figure S1**). By sequencing both the activation domain and barcode regions in unsorted cells, we filtered out barcodes attached to incorrectly synthesized mutants and use a set of high confidence barcodes for all analyses below (**Figure S2A**). The sensitivity of the assay is a 0.4-fold change in activity (p < 0.05, Bonferroni corrected two-sided t-test) and the dynamic range is >10-fold, which together provide excellent measurement resolution.

### A library of rationally designed mutants

To study how sequence-encoded physical properties control activation domain strength, we designed a library of Gcn4 activation domain mutants that systematically varies net charge, hydrophobicity, and predicted disorder (**
Figure 1B**). These mutations were designed to test several hypotheses: the *acid blob* hypothesis which predicts that net negative charge is the key property of activation domains (Ma and Ptashne, 1987a; Sigler, 1988), the *hydrophobicity* hypothesis which predicts that net hydrophobicity controls activation domain strength (Cress and Triezenberg, 1991; Jackson et al., 1996), and the *intrinsic disorder* hypothesis which predicts that structural flexibility is an essential property of activation domains (Dyson and Wright, 2016) (See Methods for design details). Hydrophobicity is calculated with the normalized Kyte-Doolittle scale (Kyte and Doolittle, 1982; Holehouse et al., 2017; Uversky, 2002) and intrinsic disorder is predicted with the IUPred algorithm (Dosztányi et al., 2005). Most mutants contain four to six substitutions and preserve the critical WxxLF motif (W120, L123 and F124) (Brzovic et al., 2011; Drysdale et al., 1995; Warfield et al., 2014). To further separate charge and disorder, we included a set of 146 mutants that convert all combinations of aspartic acids to glutamic acids (D -> E) and all combinations of glutamic acids to aspartic acids (E -> D); D and E are both negatively charged, but have different effects on predicted disorder (Oldfield and Dunker, 2014). Compared to a mutational scan that converts every position to all other amino acids (Fowler and Fields, 2014; McLaughlin et al., 2012), this rational mutagenesis covers a 3.5 fold larger range in charge (-2 to +11 compared to ±2), a 4-fold larger range of hydrophobicity and a 2.3-fold larger range of predicted disorder (**
Figure 1B**). More importantly, our rationally designed mutations change one property at time, independently testing each hypothesis.

We designed an additional set of mutants to test the hypothesis that embedding the WxxLF motif in an IDR is sufficient to create an activation domain (Warfield et al., 2014) by shuffling the 41 amino acids surrounding this motif. Each of the resulting 1224 “shuffle” mutants has a different linear sequence with the same net charge, hydrophobicity, and predicted disorder as the wild type (WT) sequence, and consequently would be located at the origin in Figure 1B. To create these mutants we randomly permuted the residues in the activation domain outside of the WxxLF motif and then selected variants that systematically changed the linear charge mixture of the sequence (Das and Pappu, 2013). Additionally, as controls we included all previously studied point mutations in this activation domain, each with 10 barcodes (Brzovic et al., 2011; Drysdale et al., 1995). Finally, we included a set of sequences that tile across the Gcn4 orthologs from 49 yeast species (Methods). In total, our library contained 6340 distinct Gcn4 mutants.

### Validating the method

We first grew our library in complete medium and quantified the activities of every library member in parallel. Our results confirm previous one-at-a-time mutant analyses (**
Figure 2A**). Mutations in the primary WxxLF motif consistently decrease activity, as shown previously (Brzovic et al., 2011; Drysdale et al., 1995; Jackson et al., 1996; Warfield et al., 2014). Some studies have also reported the presence of a secondary motif, MxxYxxL, and our results confirm that mutations in these residues (M107, Y110, and L113) also decrease activity (**
Figure 2A**) (Drysdale et al., 1995; Jackson et al., 1996). This secondary motif is enriched in the most active variants and depleted in the least active variants (p<0.01, Bonferroni corrected Fisher’s exact test, **Figure S2B**). Consistent with previous studies, mutating particular acidic residues to alanine has, on average, no effect or causes small increases in activity. The helix breaking S122P mutation causes the previously reported modest decrease in activity (Brzovic et al., 2011). Our measurements correlate well with the effects of Gcn4 mutants measured by quantitative PCR at the *ARG3* promoter (*R_P_* = 0.76 *R_s_* = 0.87) (Brzovic et al., 2011). These findings demonstrate the accuracy of our method.

**Figure 2:**
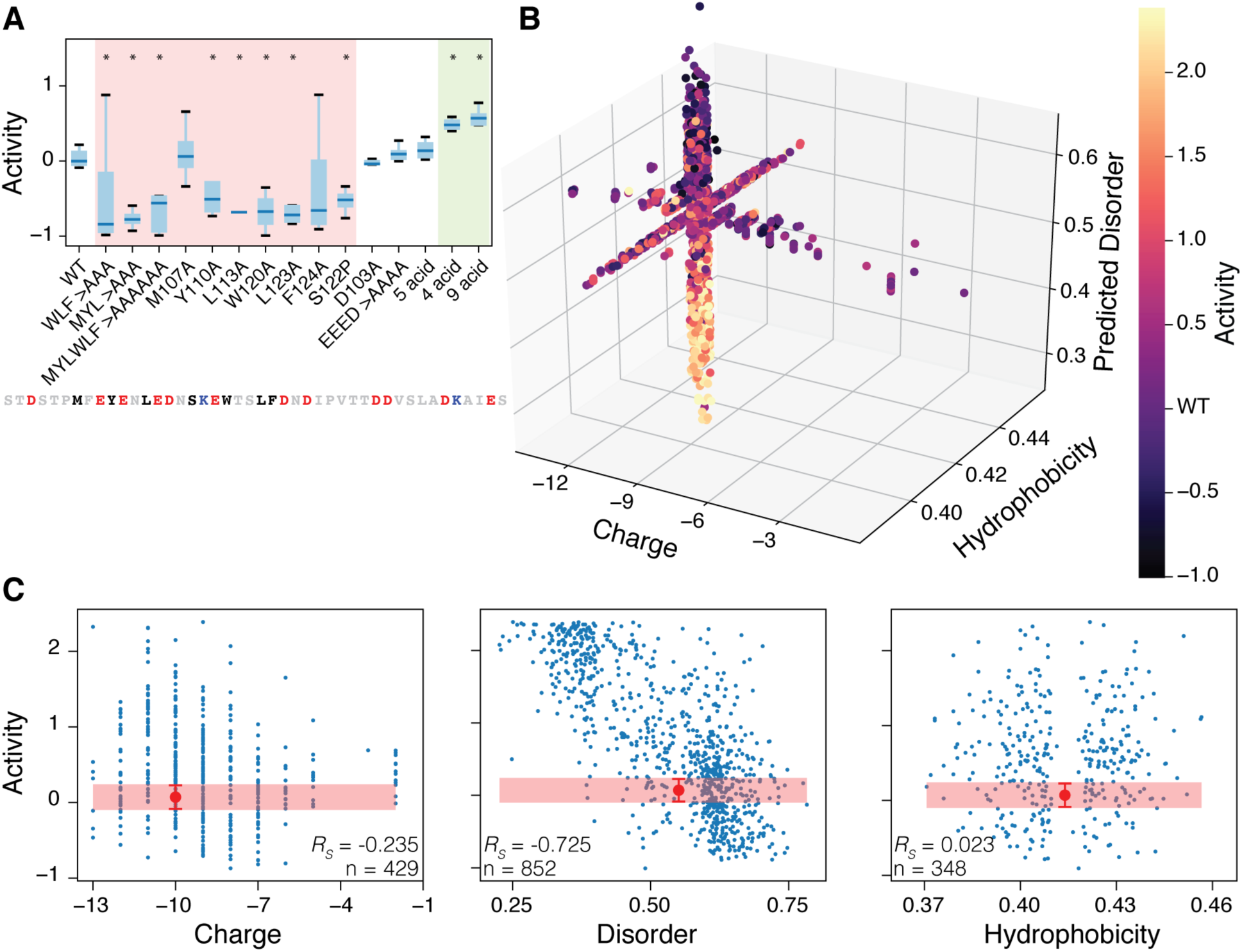
Activity measurements for the library in complete media. (**A**) High-throughput measurements match published measurements of mutations that decrease (pink), increase (green), or do not change (white) activity. In all figures, activity is log_2_(raw activity/median WT activity). (*) indicates mutants that are significantly different from WT (p<0.05, Student’s two-sided t-test). The Gcn4 central acidic activation domain sequence is shown highlighting key hydrophobic (black), acidic (red) and positively charged (blue) residues. The acidic residue mutants are named after the original works: *EEED* contains E109A, E111A, E114A, D115A (Drysdale et al., 1995); *5 acid* contains E109A, E111A, E114A, D115 E119A; *4 acid* contains D125A, D127A, D133A and D134A; *9 acid* combines *5 acid* and *4 acid* (Brzovic et al., 2011). See Figure S2B for individual data points. (**B**) Activity of 1,738 mutants from the systematic mutation sets that were measured with high confidence. Each point is plotted according to its net charge, net hydrophobicity and predicted disorder and colored according to its activity. Mutations that decrease predicted disorder increase activity. (**C**) For each set of systematic mutants, the varied property is plotted against activity. The red region and error bars indicate the mean and standard deviation of WT activity calculated from measurements from 8 barcodes.

We next examined the overall trends in the mutants designed to separate the effects of charge, hydrophobicity, and predicted disorder (**
Figure 2B**). The strongest signal was a negative correlation between predicted disorder and activity (*R_P_* = -0.76, *R_s_* = -0.73 in the disorder variants, **
Figure 2C**). Mutations predicted to make this region more ordered (i.e. lower IUpred score) also make it stronger in our assay. Net charge was not correlated with activity, as expected from the literature. Similarly, there was no correlation between net hydrophobicity and activity. However, both the charge and hydrophobicity mutants spanned the entire dynamic range in our measurements, suggesting that the effects of altering these properties are context specific, a hypothesis we explore below.

### The specific activity of the Gcn4 activation domain increases during starvation

We identified a new layer of Gcn4 induction in response to nutrient stress. Nutrient stress triggers the translation of *GCN4* mRNAs, and the resulting Gcn4 protein is a critical stress response TF that activates amino acid biogenesis genes (Hinnebusch, 2005; Niederberger et al., 1981). This role for Gcn4 prompted us to assay the library during amino acid starvation. Specifically, we washed cells growing in synthetic complete media and resuspended them in media without amino acids. When we compared the measurements made in complete medium to amino acid starvation conditions we found that the specific activity of the wild-type (WT) Gcn4 activation domain increased by 64%. We define *induction* as the ratio of specific activity during starvation to the specific activity in complete media (i.e. the fold change in activity, WT= 1.64). This induction occurs in our synthetic TF system, and must be independent of the normal Gcn4 translational response to stress (Hinnebusch, 2005). Our data indicate that starvation induces an increase in the specific activity of the central acidic activation domain of Gcn4 on top of the well-established translational induction of the protein (Hinnebusch, 2005).

Compared to most mutants, the WT protein has lower baseline activity, but is more inducible during starvation. Most mutants (68%) have higher activity than the WT in complete media. However, during amino acid starvation, only 8.6% of mutants are more induced than WT (**
Figure 3**, Figure S3A,B). Some mutants fail to induce because they are already at maximum levels of activity, but most mutants with activity levels similar to wild-type in complete media fail to induce or have little induction during amino acid starvation. In fact, most mutants in our collection were less inducible than the WT protein. These data may explain why the Gcn4 activation domain is weak; selection may have favored inducibility over activity.

**Figure 3:**
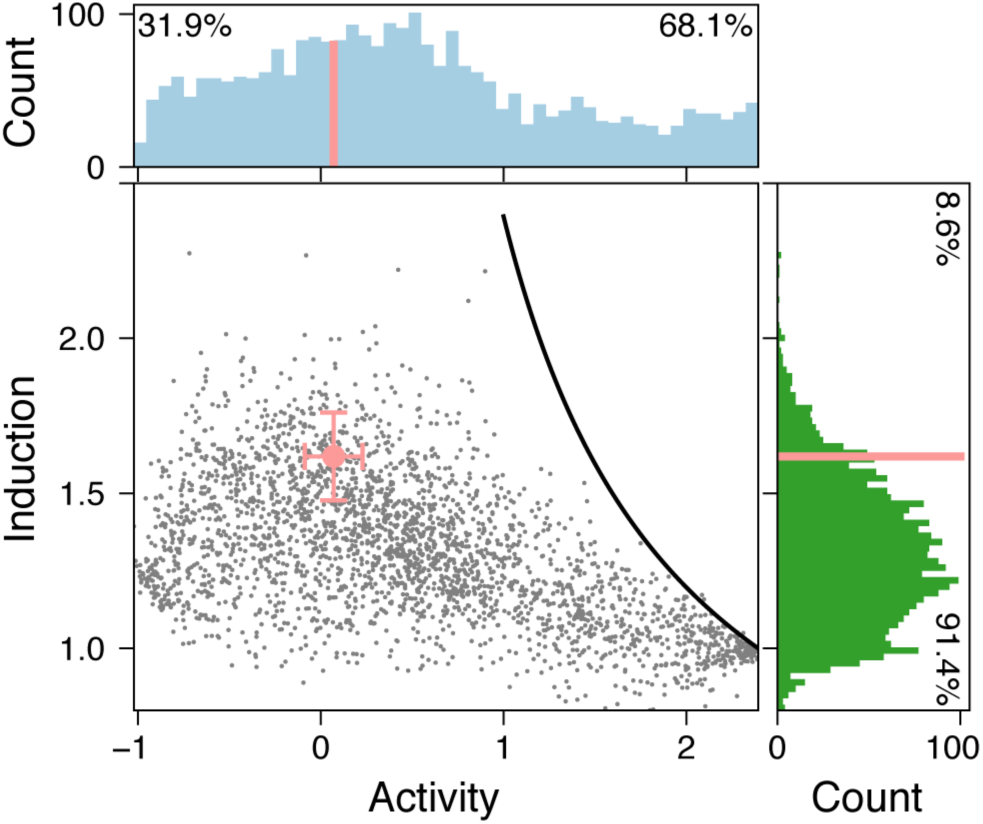
The WT Gcn4 activation domain is less active and more inducible than most mutants. Each variant is plotted by its activity in complete media and its induction, the fold change in activity during starvation. The detection limit for induction is shown in black. Histograms of activity in complete media (light blue, top) and induction (dark green, right) reveal that the WT Gcn4 activation domain (pink) is not very strong but is highly induced. Error bars are mean and standard deviation. See Figure S3 for raw changes in activity. N = 2716 from the high confidence data set.

### All atom simulations probe the predicted changes in disorder

To probe for a structural explanation for the observed negative correlation between predicted disorder and activity, we performed all-atom Monte Carlo simulations (Das et al.,2016; Martin et al., 2016; Vitalis and Pappu, 2009). These simulations generate conformational ensembles of IDRs that provide insight into the coupling between amino acid sequences and conformational behavior, and have provided extensive insight into a wide range of intrinsically disordered regions (Das et al., 2016; Fuertes et al., 2017; Martin et al., 2016; Wei et al., 2017). We applied this technique at unprecedented scale by examining over 400 distinct sequences from our library with ten independent simulations each.

We explored three explanations for the negative correlation between predicted intrinsic disorder and activity. First, the increase in order could be driven by increased secondary structure. This hypothesis is based on the observation that the WxxLF motif is unstructured in solution, but forms an α-helix when bound to the coactivator Gal11/Med15 (Brzovic et al., 2011), and preformation of an α-helix can increase activity (Warfield et al., 2014). Second, the decrease in disorder could be driven by a collapse of the polypeptide into a semi-folded state that resembles the dense cores of globular proteins in the IUpred training set. Third, the correlation could result from a hidden covariate in the prediction algorithm. We tested the first two possibilities with simulations and the third by statistical analysis.

First, we examined the idea that the increase in predicted order could be driven by increased secondary structure. Contrary to this expectation, when we simulated the fifty least and fifty most active mutants from the disorder set, there is no significant difference in the frequency of mutants predicted to form helices (Figure SS4A). Next, we tested the explanation that the predicted disorder is due to the collapse of the peptide chain. The simulations showed that most active mutants have a smaller radius of gyration (10% smaller, bootstrap *p* < 1×10^-5^ Figure S4B) than inactive mutants, supporting this possibility. The simulations do not indicate the large changes in intrinsic disorder predicted from the IUPred algorithm used to design the mutants (Dosztányi et al., 2005). Highly active variants adopt somewhat more compact conformations yet remain intrinsically disordered. The simulations suggest that causing large changes in the intrinsic disorder of the Gcn4 activation domain with substitutions is difficult, perhaps because the sequence favors disordered conformations.

Finally, we considered the possibility that the IUPred algorithm predicted the activity of Gcn4 mutants because of a hidden covariate. In an ANOVA analysis, the predicted disorder score explains less of the variance in activity (57.2%) than simply counting the number of W, F, Y, M, and L residues (66.5%) (Table S1). Adding predicted disorder to the counts of these five amino acids provides very little added explanatory power (67.1%). Closer inspection of the IUPred algorithm revealed that these five amino acids contribute heavily to the disorder prediction (Dosztányi et al., 2005). We speculate that the IUPred algorithm predicts changes in the strength of mutants in part because it counts the right residues.

### Simulations reveal that exposure of aromatic residues predicts activity

When we searched the simulation data for structural features that predict activity, the most and least active mutants are separated by the ratio of solvent accessibility of aromatic residues (W, Y and F) to radius of gyration (disorder variants, **
Figure 4A**; all variants, Figure S4E). The most active variants have more aromatic residues than the least active, but the increased solvent exposure of these residues is not determined only by their abundance: when we simulated another well-studied intrinsically disordered region from yeast, adding extra aromatic residues did not increase overall solvent exposure because these aromatic residues drive compaction, resulting in little net change in solvent accessibility (Methods, Figure S4D). Variants that can adopt compact ensembles of conformations while keeping their aromatic residues exposed to the solvent are most active in our assay.

**Figure 4:**
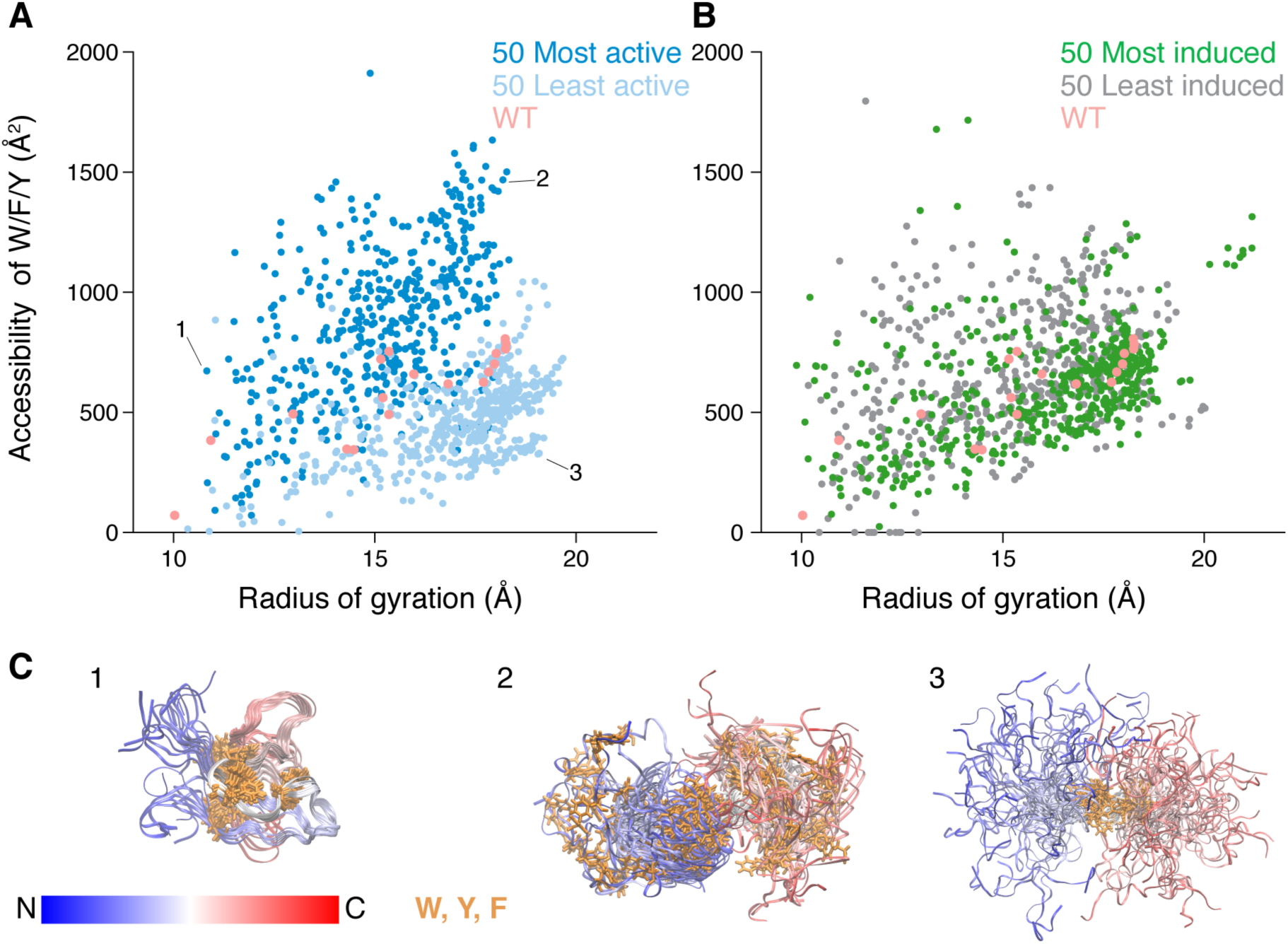
All atom simulations reveal that highly active variants keep aromatic residues exposed to solvent. (**A**) From the variants that perturbed disorder, we simulated the 50 most active (dark blue) and least active (light blue) variants. For each mutant, we performed ten independent simulations, and plotted each simulation by the radius of gyration and accessibility of aromatic residues. Highly active variants have a higher ratio of aromatic accessibility to radius of gyration compared to lowly active variants. WT (pink) sits between the populations. (**B**) The 50 most induced variants under amino acid starvation (dark green) cluster around WT. The 50 least induced variants (light green) were selected to span the same range of expression as the most induced set, and correspondingly show similar aromatic exposure and radius of gyration. (**C**) For the 3 simulations indicated in A, the final 50 snapshots are overlaid and aligned on the central region. The peptide backbone is colored from N (blue) to C (red) terminus and the aromatic residue sidechains are colored orange. In the first panel, the backbone is collapsed in a way that leaves a phenylalanine on the surface. In the second panel, backbone is extended, exposing the distal aromatic residues. In the third panel, the backbone is expanded but all the aromatic residues are sequestered in the center.

Simulations of the WT Gcn4 activation domain reveal an intermediate radius of gyration and aromatic exposure, consistent with its intermediate activity. Compared to all other intrinsically disordered regions in the yeast proteome, the Gcn4 activation domain is highly acidic (93.5^th^ percentile in net negative charge per residue) and highly hydrophobic (93^rd^ percentile in the percentage of Y/W/F/L/I/V/M residues). In this activation domain, the tendency of highly hydrophobic side chains to drive collapse appears to be balanced by the tendency of strongly acidic regions to resist collapse due to electrostatic repulsion (Müller-Späth et al., 2010). Taken together, our results suggest that the sequence of the Gcn4 activation domain makes it particularly adept at exposing aromatic residues to solvent, which may facilitate protein-protein interactions. If the key to activity is aromatic exposure, the α-helix may merely be a convenient, but not necessary, way to display these residues to coactivators such as Gal11 (Brzovic et al., 2011). This possibility would explain why we do not observe an increase in helical content in the highly active mutants.

In additional simulations, the 50 most inducible and least inducible variants had aromatic accessibility and radii of gyration similar to WT, in between the high and low activity mutants (**
Figure 4B**). The highly inducible variants require some aromatic exposure to be active, but additional exposure does not boost inducibility.

### Aromatic residues control activity

While there is no relationship between net hydrophobicity and activity (or inducibility), particular amino acids do have an effect. Adding aromatic residues (W, F, and Y) increases activity in complete media (**
Figure 5A,B,C**, Figure S5) as seen previously (Warfield et al., 2014). Adding or subtracting aromatic residues decreases inducibility (**
Figure 5E,F,G**, Figure S6). Given the long literature on the importance of hydrophobic residues controlling activation domain strength (Brzovic et al., 2011; Cress and Triezenberg, 1991; Drysdale et al., 1995; Jackson et al., 1996; Lu et al., 2005; Lum et al., 2006; Warfield et al., 2014), we emphasize that adding the very hydrophobic residues Isoleucine (I) and Valine (V) does not boost activity, and that Methionine (M) and Leucine (L) have only modest activating effects (Figure S5). Interestingly, the boost in activity from the addition of an aromatic depends on the context. For example, adding a W near the primary WxxLF motif causes a greater-than-average increase in activity, while adding a W in the secondary MxxYxxL motif causes a smaller-than-average increase in activity (**
Figure 6A**). Adding an aromatic residue almost always increases activity, but its location modulates this increase in a predictable way. To first approximation, the number of aromatics residues controls the intrinsic activity of mutant activation domains.

**Figure 5:**
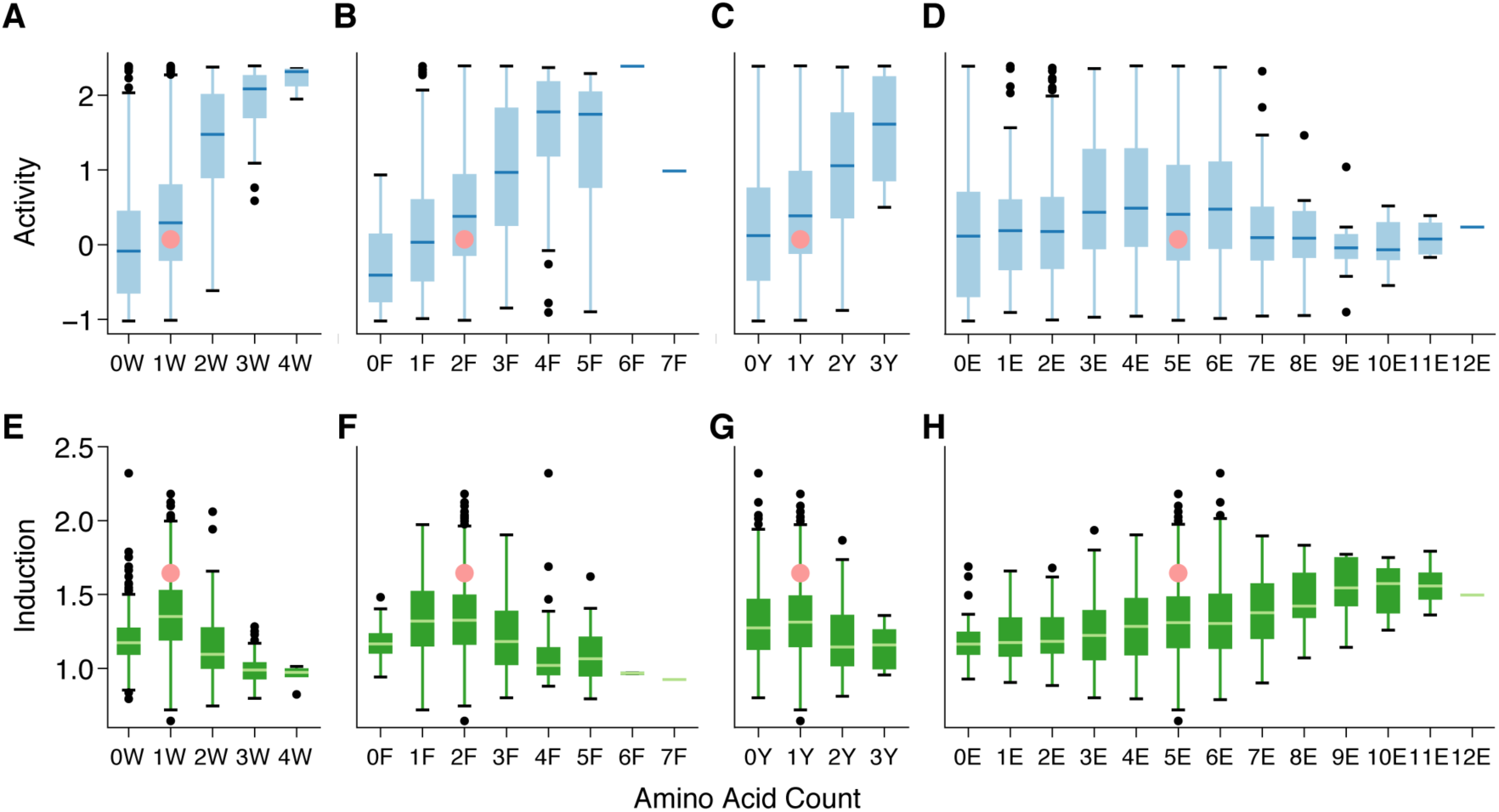
Aromatic residues control activity. Box plots show the activity (blue) of all variants in the library containing the indicated number of tryptophan (W), phenylalanine (F), tyrosine (Y) or glutamic acid (E) residues. Aromatic residues increase activity (blue) in complete media (**A-C**) but have a mixed effect on induction (green) during amino acid starvation (**E-G**). WT (pink) has low activity, but has high inducibility. Glutamic acids (E) have little effect on activity (**D**) but increase inducibility (**H**). N = 2716.

**Figure 6:**
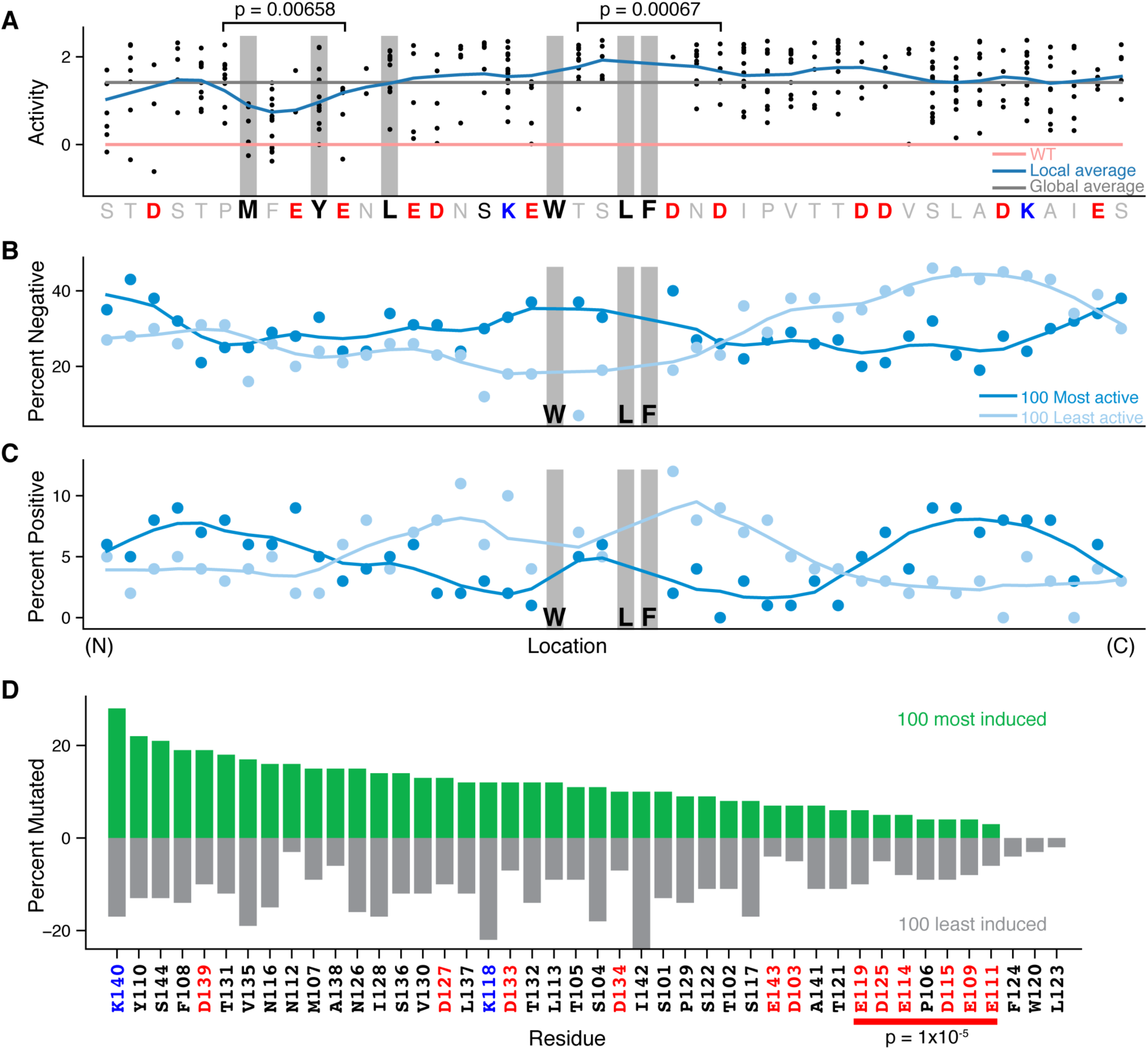
Local sequence context modulates the effects of a mutation. (**A**) Each dot indicates the activity of a variant that adds a tryptophan (W) at that position. Data are from the 310 charge, hydrophobicity and disorder variants with exactly two Ws. Adding a W near the WxxLF motif gives a larger boost in activity than adding a W near the MxxYxxL motif. The divergence of local average (dark blue; lowess smoothing) from the global average (dark gray) indicate context specific effects. Most variants that add a W are stronger than WT (pink). Brackets indicate five consecutive positions that have a local mean significantly different from the global mean in permutation tests. (**B**) For the 100 most active (dark blue) and 100 least active (light blue) shuffle variants, the fraction of sequences that have a negatively charged residue at each position is plotted along with a smoothed line (lowess). (**C**) As in B, but for positively charged residues. (N) and (C) denote the N and C termini, respectively. Note only the WxxLF positions are preserved in the shuffle mutants. (**D**) The number of times each position is mutated in the 100 most induced variants (dark green) and the number of times each position is mutated in the 100 least induced variants (light green). This analysis excludes the shuffle mutants and orthologs. Acidic (negatively charged) residues are colored in red and positively charged residues are colored in blue. No individual position is mutated more than expected by chance (p<0.01 Fisher’s Exact Test, Bonferroni corrected), buthe six central acidic positions (red bar) are under mutated (p=1×10^-5^ in permutation tests).

### Acidic residues create a permissive context for the WxxLF motif

The role of acidic residues in acidic activation domains has long been debated (Brzovic et al., 2011; Cress and Triezenberg, 1991; Drysdale et al., 1995; Ma and Ptashne, 1987a; Sigler, 1988; Warfield et al., 2014). We find that overall net charge is not predictive of activity, but the local distribution of charge has a large impact on activity. This effect is most visible in the shuffle variants. While all the shuffle mutants have the WT number of acidic residues, highly active variants accumulate negatively charged residues near the WxxLF motif, while inactive variants accumulate negatively charged residues on the C-terminal end and positively charged residues in the center (**
Figure 6B,C**, Figure S7A). In the full set of shuffle mutants, local net negative charge is most correlated with activity around the WxxLF motif (Figure S7B). This result makes sense in light of the NMR structure of the Gcn4 activation domain bound to Gal11 (Brzovic et al., 2011), because the hydrophobic binding cleft on Gal11 where the WxxLF motif docks is flanked by positively charged residues. These observations are consistent with electrostatic interactions guiding Gcn4-Gal11 binding, as has been observed in other disordered regions that undergo coupled folding and binding (Rogers et al., 2013). Contrary to previous predictions, the shuffle mutants demonstrate that the WxxLF motif is not sufficient to convert any disordered peptide into an activation domain. Instead, this motif requires an appropriate negatively charged context. Together with the simulations, our data support a model where negative charges keep hydrophobic residues exposed to solvent where they can interact with binding partners.

### Acidic residues control inducibility

The central acidic residues in the Gcn4 activation domain are important for inducibility. The acidic residues around the primary and secondary motifs are six of the seven least mutated positions in the 100 most inducible variants (**
Figure 6D**). When considered alone, none of these positions is mutated less than expected by chance (Fisher’s Exact Test), but collectively they are significantly under mutated (*p* = 1×10^-5^ permutation test). Mutations that changed acidic residues to alanine decrease inducibility (Figure S3D). We do not see evidence for phosphorylation being involved in induction: T105 is the key site of phosphorylation in the starvation responsive degron (Kornitzer et al., 1994), but this position is not mutated more often in either the most or least induced variants (**
Figure 6D**), nor in the most or least stable variants. Across all of our mutants, the number of glutamic acids has little relationship with activity in complete media (**
Figure 5D**), but is positively correlated with induction (**
Figure 5H**). However, very few of these mutants are as inducible as WT, emphasizing the importance of the local placement of acidic residues. Together with the activity data from the shuffle mutants, these data demonstrate, for the first time, a functional role for acidic residues in an acidic activation domain.

### A gene specific model for how mutations impact function of the Gcn4 central acidic activation domain

We used physical properties and amino acid composition to train support vector regression models to predict the impact of mutations in the Gcn4 activation domain. We excluded the shuffle mutants and the orthologs from this analysis. After exploring a large family of models, the final model has six parameters: the number of W’s, F’s, Y’s, L’s and M’s in the full sequence and the net charge in the central region (positions 112-125). This central charge effect is consistent with the charge effect seen above for the shuffle mutants, even though those mutants were not included in the machine learning analysis. Under five-fold cross validation, this regression explains 58% of the variance in activity, enough to be a useful predictor (**
Figure 7A**). Overall the machine learning is consisent with the primary trends in the data: the number of key hydrophobic residues, primarily aromatics, and charge in the central region control Gcn4 activation domain activity.

**Figure 7:**
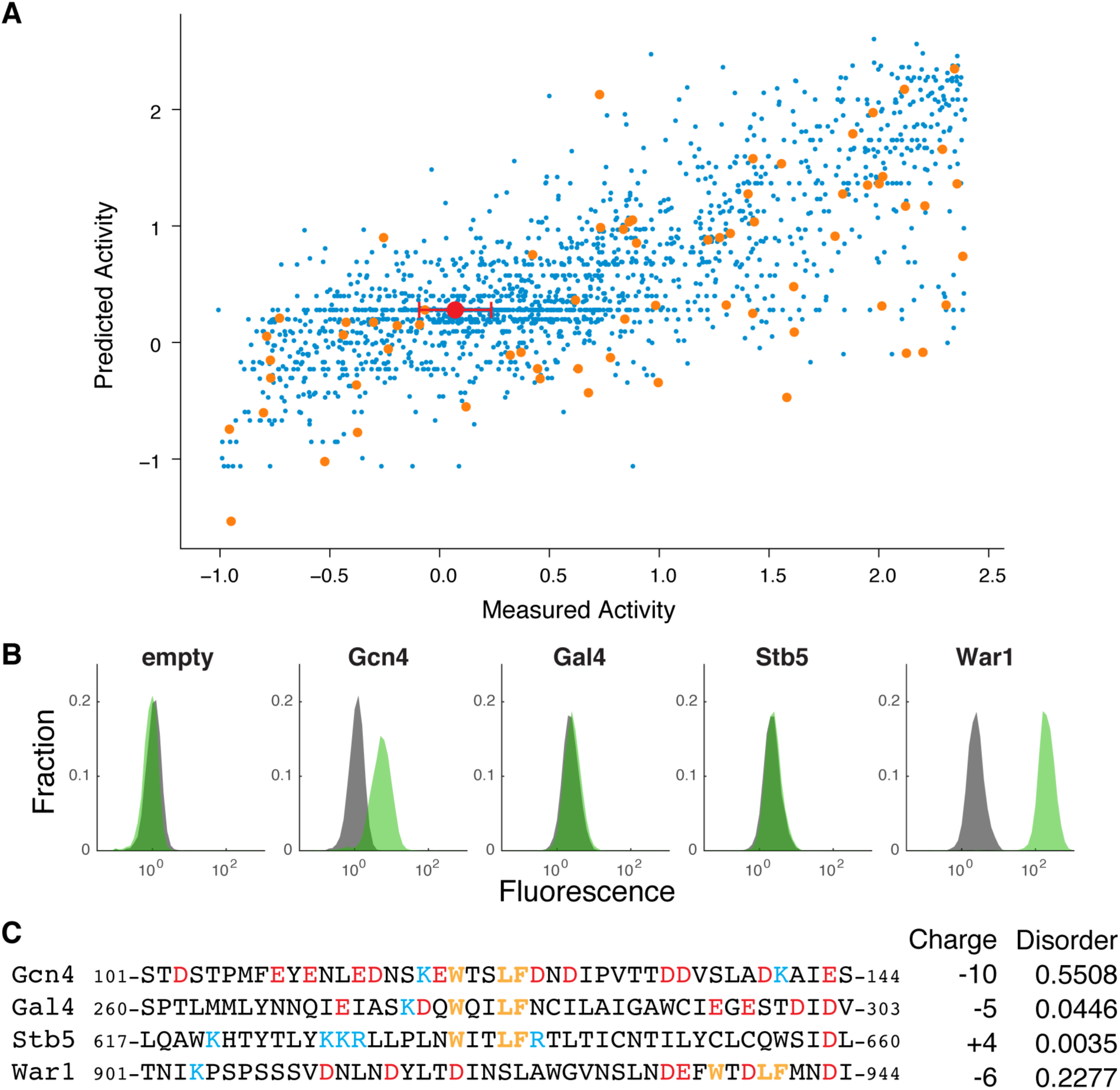
Predicting activation domains in the proteome. (**A**) We created a gene specific model for how mutations control the activity of the Gcn4 central acidic activation domain by training a support vector regression model on 1738 substitution mutants. The WT mean and standard deviation is shown in red. When the model is trained on all the data and used to predict all the data (shown), the R^2^ is 0.59. After 5-fold cross validation, the average R^2^ is 0.58. We further tested the model with the direct orthologous sequences (orange), which were not included in the training data. (**B**) For each of the four TFs in yeast that contain both a DNA binding domain and a WxxLF motif, we individually tested if this region has activation domain activity. For each construct, the distribution of fluorescence values from the GFP reporter in the presence (green) or absence (grey) of ß-estradiol. ß-estradiol induces nuclear localization of the synthetic TF. If the attached sequence lacks activation domain activity, these distributions overlap; when an active activation domain is attached, nuclear localization induces expression of the GFP reporter. (**C**) In the protein sequences, the WxxLF motif (orange) negatively charged residues (red) and positively charged (blue) residues are highlighted. Net charge and predicted intrinsic disorder are shown.

### Predicting activation domains in the proteome

As a test of our unified model that acidic residues create a permissive context for hydrophobic motifs, we examined whether other instances of the WxxLF motif in the yeast genome are part of activation domains. The WxxLF motif occurs in three other TFs: Gal4, Stb5 and War1. The region from Gal4 is known to not be an activation domain (Ma and Ptashne, 1987b; Warfield et al., 2014). In Gal4 and Stb5 the motif occurs in regions with low predicted disorder and without nearby acidic residues, while the War1 region is predicted to be disordered and has negative charges around the motif. Based on these context features we predicted that only the War1 region would be active in our assay. We tested 44 amino acid regions centered on the motif (for War1 we tested the final 44 residues of the protein). Consistent with our predictions, only the War1 WxxLF region functions as an activation domain (**
Figure 7B**). We conclude our model predicts active instances of the WxxLF motif in the proteome. This result is a step towards combining motifs and context features to predict activation domains.

We could not apply our SVR predictor more generally to the yeast proteome because the model is trained specifically on Gcn4 variants with the WxxLF motif. A model trained on a more diverse set of inactive sequences might have more power to detect other classes of acidic activation domains. Instead we considered an alternate approach to identifying additional candidate activation domains in the yeast proteome. As noted above, compared to other disordered regions in the proteome, the Gcn4 activation domain is highly acidic (93.5th percentile in net negative charge per residue) and highly hydrophobic (93rd percentile in the percentage of Y/W/F/L/I/V/M residues). Based on these properties we speculate that disordered regions with extreme values for both acidity and hydrophobicity are good candidates for additional activation domains. We scanned all 44 amino acid windows in the proteome for three criteria: IUpred > 0.4, net negative charge > Gcn4, and Y/W/F/L/I/V/M count > Gcn4. We found 1004 windows (0.12% of all windows tested) on 204 proteins that met these criteria (Supplemental Table 4). Half of these proteins are localized to the nucleus according to GO enrichment. These windows may be enriched for activation domains or protein-protein interaction regions. When we repeated this analysis on 163 proteins annotated to contain DNA binding domains, we found 109 windows (0.12%) that overlap to create 11 regions on 11 proteins. These disordered sequences, which balance acidity and hydrophobicity similarly to Gcn4, may be enriched for activation domains.

## Discussion

Why are activation domains acidic? Here we show that acidic residues make the Gcn4 activation domain inducible during amino acid starvation. This role for acidic residues explains the long standing puzzle of why this activation domain is weak, and why many mutations can make a synthetic activation domain with the WxxLF motif much stronger (Drysdale et al., 1995; Warfield et al., 2014). We speculate that the Gcn4 central acidic activation domain is under evolutionary pressure to be weak but inducible.

The most likely molecular mechanism for inducibility is a change in the availability (or activity) of a coactivator during starvation. The inducibility of Gcn4 may reduce the fitness cost of leaky translation during non-starvation conditions (Drysdale et al., 1995; Warfield et al., 2014), may endow the circuit with a greater dynamic range, or enable the starvation response to sense fold changes in nutrient availability (Goentoro et al., 2009).

Hydrophobicity and acidity control activation domain activity in contrasting ways. The number of aromatics coarsely sets activity in multiple conditions while the acidic residues can tune activity between conditions, controlling inducibility. Acidity further creates a permissive context for hydrophobic residues to function. Classic mutations that change negatively charged residues to alanine make the peptide much more ordered and likely keep the aromatics exposed to coactivators, explaining why these mutations can sometimes increase activity. A molecular explanation for the contrasting roles for aromatic and charged residues may lie in the biophysics of activation domain coactivator binding. The binding kinetics of some acidic activation domains (Gal4 and VP16) are biphasic, with a fast, charge-mediated step followed by a slower, high-affinity hydrophobicity-mediated step (Ferreira et al., 2005). The complementary effects of hydrophobicity and charge may therefore arise through their influence on different aspects of the binding kinetics of Gcn4 with its coactivators.

Our data support a model in which acidic residues keep key hydrophobic residues exposed to solvent where they can interact with coactivators. There are hints of this model in the literature (Cress and Triezenberg, 1991; Lu et al., 2005; Lum et al., 2006; Ruden, 1992; Shen et al., 1996a, 1996b; Sullivan et al., 1998). This model explains why acidic activation domains are acidic, and explains why mutating hydrophobic residues has the largest impact on activity. In this model, α-helices are a convenient, but not essential means to expose key hydrophobic residues; explaining why inserting prolines into α-helices only moderately impairs activity in many activation domains (Brzovic et al., 2011; Cress and Triezenberg, 1991). Intrinsic disorder, particularly the ability of prolines to cause expansion (Martin et al., 2016), may be another means of keeping aromatic residues exposed to coactivators in proline-rich activation domains.

The potential for a single activation domain sequence to have different intrinsic activities in different cellular states could be a useful feature for tuning gene regulation in development. Different cell types deploy specialized paralogs for some components of Mediator and the basal transcriptional machinery (Goodrich and Tjian, 2010; Malik and Roeder, 2010). If an activation domain has a stronger affinity for a particular paralog that activation domain will be most active in a limited subset of cell types. The potential for activation domain activity to respond to cellular context may be a general feature of cell type specific gene regulation.

## METHODS

### Yeast strains and media

FY4, is a MATa prototroph and a gift from Fred Winston (Cohen Strain 1091, MY8). FY5 is a MATalpha prototroph and a gift from Fred Winston (Cohen Strain 1092, MY9). Mating types were confirmed by colony PCR. Both strains grow on minimal media, confirming they are prototrophs.

Strains were grown in synthetic complete (SC) dextrose media at 30^o^C (Amberg et al., 2005). Overnight cultures were diluted (1:5) into SC+1 uM ß-estradiol and grown for 3.5-4 hours at 30^o^C before FACS. For the amino acid starvation experiments, we washed the overnight culture with water twice and resuspened in minimal media without amino acids (synthetic dextrose, SD) with 1 uM B-estradiol for 4 hours before FACS. YDP plates and media were used for transformations (Amberg et al., 2005).

### Design of Gcn4 Activation Domain Mutants

We designed the systematic charge, hydrophobicity and disorder variants by generating random mutations at 4-6 positions, calculating the properties of each variant with localCIDER (Holehouse et al., 2017) and keeping candidates which changed only one parameter. To calculate net hydrophobicity we used the normalized Kyte Doolittle scale (Kyte and Doolittle, 1982; Uversky, 2002). We predicted intrinsic disorder with a custom python implementation (A. S. H.) of the IUPred algorithm (Dosztányi et al., 2005). In most cases we protected W120, L123 and F124 from mutation. To probe a wider range of charge variants, we added variants that converted all combinations of charged residues to the opposite charge or neutral residues (size matched D->N, E->Q). In addition, we converted polar residues (S and T) to charged residues and filtered for variants with small changes in disorder and hydrophobicity. We supplemented the disorder variants with a set of 146 mutations that interchanged D and E, which we call the HoldChargeDisorder variants. For the 5 E’s we mutated 1-5 positions to D. For the 7 D’s, we mutated 1-7 to E and removed duplicates. To generate the shuffle mutants, we permuted all residues except W120, L123 and F124, computed a charge mixture statistic (*κ*, as described in(Das and Pappu, 2013)) and used the Wang-Landau algorithm to flatten the distribution (Wang and Landau, 2001). These permutations do not change net charge and net hydrophobicity but caused small changes in intrinsic disorder, so we filtered for variants that changed predicted disorder by less than 1%.

In our mutation selection scheme, adding tryptophan (W) causes a large change in disorder and only a small change in hydrophobicity, making it easier for a second mutation to compensate for the increased hydrophobicity. In the normalized Kyte Doolittle hydrophobicity scale (Kyte and Doolittle, 1982; Uversky, 2002), W is not very hydrophobic compared to I or V. Consequently, the disorder variants show the wider range in the number of Ws than the systematic hydrophobicity mutants.

As controls, we included 15 published sequences. The EEED >4xA construct contains E109A, E111A, E114A and D115A (Drysdale et al., 1995). The ‘5 acid’ construct contains E109A, E111A, E114A, D115A and E119A (Brzovic et al., 2011). The ‘4 acid’ construct contains D125A, D127A, D133A and D134A (Brzovic et al., 2011). The ‘9 acid’ construct combines the mutations in the ‘5 acid’ and ‘4 acid’ constructs (Brzovic et al., 2011). Each control sequence was included in the library 10 times with 10 unique barcodes. The WT amino acid sequence was included with five different codon usages and 23 barcodes. All other variants had one barcode. Additionally, we systematically moved the WxxLF motif across the Gcn4p activation domain in three ways: we moved WXXLF, WTSLF and KEWTSLFDN to every position in the peptide. We also added WTSWF, WTSLW, WTSWW and WSTLF at the native position. We did not recover enough of these ‘move motif’ mutants (27/117) to make any reliable conclusions.

For the direct orthologs, we used MUSCLE to create an alignment from 49 species (Edgar, 2004).The ortholog list (Supplemental Data Table 3) was curated by hand from the Yeast Gene Order Browser, the Broad Orthogroup Repository and Fungal BLAST tool on SGD (Byrne and Wolfe, 2005; Wapinski et al., 2007). For all orthologous regions less than 44 residues, we extended them to 44. For all orthologous regions longer than 44 residues, we included two overlapping fragments, the first 44 and the final 44 residues. In addition, we tiled across each ortholog with 44 amino acid windows spaced every 22 residues. We did not recover enough of these tiling fragments to make any strong conclusions about the orthologs. In all, we designed 6500 constructs.

### The synthetic TF and reporter construct

The reporter plasmid (pMVS102, BAC1017) contains the P3 promoter (McIsaac et al., 2013b) driving a fast maturing GFP variant (Cormack et al., 1996; Iizuka et al., 2011) in a ClonNAT marked pFA6 plasmid(Longtine et al., 1998). We PCR amplified GFP from me118 (Sherman et al., 2015) (CP17.P3 CP17.P4), synthesized the the P3 promoter described in McIsaac et al. 2013a (gBlock, IDT) and added them to a ClonNAT marked pFA6 plasmid (pMVS008, MSP11 a gift from Michael Springer) digested with BamHI-HF and AscI with Gibson assembly (NEB E2611L).

The TF chassis plasmid (pMVS142, BAC1027) contains the yeast *ACT1* promoter (PCR amplified from MSP66, a gift from Michael Springer), Mouse Zif269 DnA binding domain (DbD; synthesized from IDT following the sequenced published in (McIsaac et al., 2013a)), the estrogen binding domain (EBD; PCR amplified from pRB3311, a gift from Scott McIsaac and David Botstein), a custom multiple cloning site (MCS NheI, PacI, AscI; synthesized from IDT), and the KANMX cassette in the pFA6 plasmid backbone (Longtine et al., 1998). In principle, the TF chassis is only slightly modified from the validated system described in (McIsaac et al., 2013a)—we added an mCherry at the N-terminus after the *ACT 1* promoter and replaced the VP16 activation domain with the activation domain library. In practice, the final plasmid was constructed via a series of intermediates by Gibson assembly. All enzymes were from NEB unless otherwise noted.

The two final plasmids, the reporter and the TF chassis, were deposited in Addgene (99048 and 99049).

pMVS142 was created from the following intermediate plasmids.

pMVS103: A PCR product with the ERV14 promoter (CP17.P5 and CP17.P6) from pMV003 (pJHK43, a gift from John Koschwanez) and a gblock (containing the Zif268 DBD, mTagBFP2 (copied from pFA6a-link-yomTagBFP2-SpHis5 AddGene Plasmid #44839), NheI and PacI; IDT) were assembled into pMVS011 (MSP23, a KAN marked variant of pFA6, a gift from Michael Springer) digested with SalI-HF and AscI. This assembly created the multiple cloning site site (MCS), NheI, PacI, AscI that was later used for library cloning.

pMVS111: The EBD was PCR amplified from pMVS025 (pRB3311) with primers CP18.P15 and CP18.P16 and assembled into pMVS103 cut with BamHI.

pMVS113: The *ACT1* promoter was amplified from pMVS004 (pJHK43, a gift from John Koschwanez) with primers CP19.P3 and CP19.P4 and assembled into pMVS103 digested with SalI-HF and KpnI-HF.

pMVS116: The estrogen binding domain (EBD) was PCR amplified from pMVS111 with primers CP21.P7 and CP21.P8, the ACT1 promoter and Zif268 DBD were amplified from pMVS113 with primers CP19.P3 and CP21.P11 and assembled into pMVS103 digested with SalI-HF and NheI-HF.

pMVS117: The estrogen binding domain (EBD) was PCR amplified from pMV111 with primers CP21.P7 and CP21.P9, the ACT1 promoter and Zif268 DBD were amplified from pMVS113 (CP19.P3 and CP21.P11), mCherry was PCRed from pMVS011 (MSP66 from Michael Springer; CP21.P10 and CP21.P5) and assembled into pMVS103 digested with SalI-HF and NheI-HF.

We created the final plasmid, pMVS142 by PCR amplifying mCherry from pMVS117 with primers CP25.P1 CP25.P2 and assembling into pMVS116 digested with KpnI.

The additional regions with WxxLF motifs in Figure 7 were ordered as gBlocks from IDT with 15 bp overhands and cloned into the NheI and AscI sites of pMVS142 by HiFi assembly (NEB).

The entire promoter and TF region of all plasmids was Sanger sequenced.

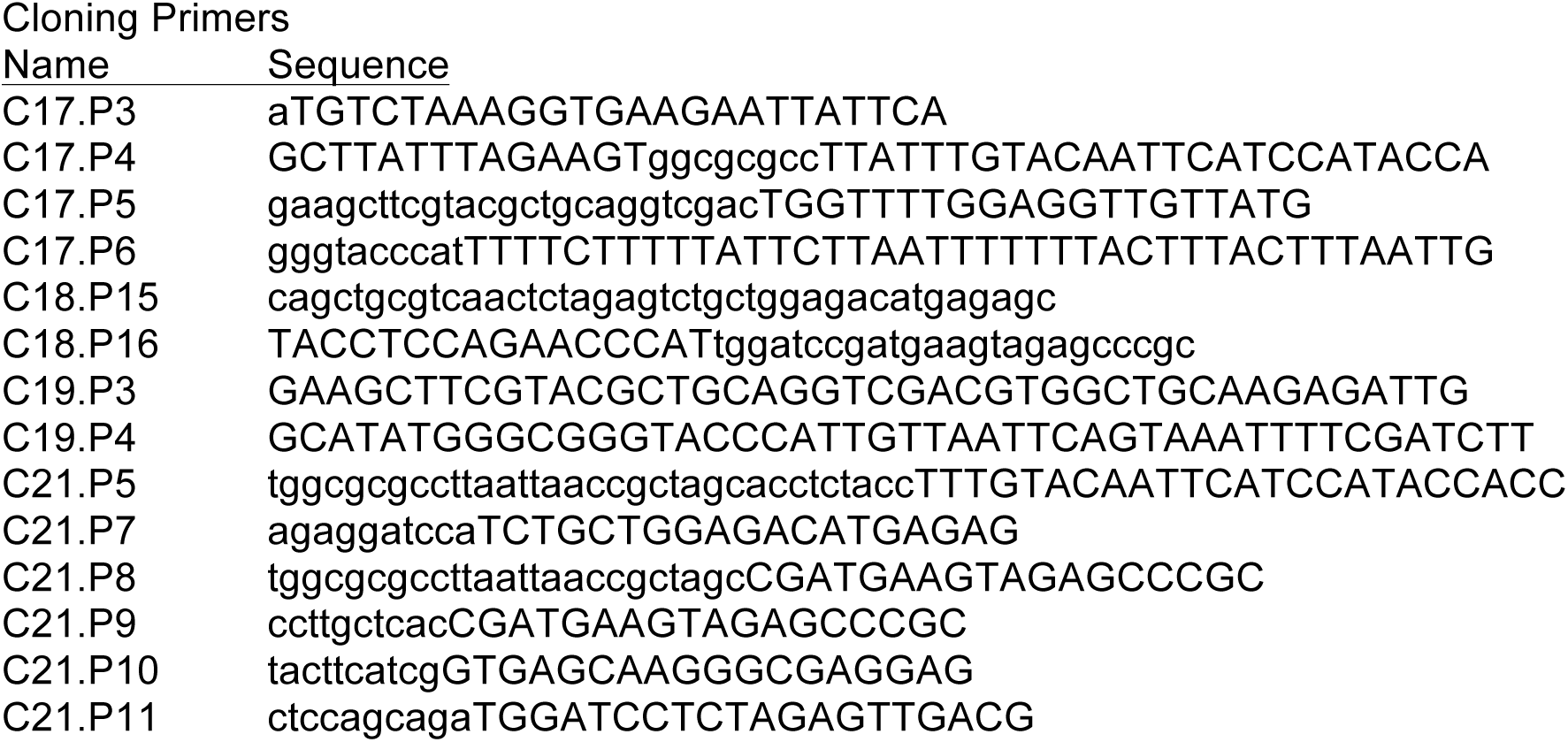

### Plasmid Library construction

We ordered a 6.5K library of 200 bp oligos from Agilent. These oligos contained a 19 bp primer (GCAGCATCTGTCC<u>GCTAGC</u>) including the NheI site, 132 bp encoding the activation domain variant, a 21 bp primer (TGATAACTAGCTGA<u>GGGCCC</u>G) with four stop codons in 3 frames and the ApaI site, a 9 bp barcode and a 19 bp primer including the AscI site (<u>GGCGCGCC</u>CTAATCTTACG). The barcodes were reused from prior work (Kwasnieski et al., 2014) and had beeen designed to have a hamming distance >2 between each sequence.

We cloned the TF library similarly to previous work in our group (Kwasnieski et al., 2012). Specifically, we used 0.5 picomoles of template and 4 rounds of PCR in 8 parallel 50 ul reactions (NEB Phusion Tm=55C with 4% DMSO), followed by PAGE on an 8% acrylamide gel. DNA was extracted from the acrylamide with dialysis cartridges (Thermo #87717) and purified by ethanol precipitation. The activation domain variants were inserted into NheI and AscI sites of the chassis plasmid (pMVS142). The backbone was digested in 16 parallel 20 ul reactions and phosphatase treated. We performed 8 parallel ligations of 20 ul for 2 hours at ~1:1 plasmid to insert ratio (NEB T4 ligase). Following electroporation in DH5a cells (NEB C2989K), we recovered ~54,000 colonies.

### Yeast library construction

To ensure only one construct per cell, we integrated our library at the URA3 locus of FY4. We used a standard three piece yeast transformation (Gietz and Woods, 2002) with 30 min at 30°C and 60 min at 42°C. To avoid PCR errors, we digested the plasmid library with SalI and EcoRI, which liberated the region spanning the *ACT1* promoter, the TF and the KANMX marker. We simultaneously digested with PacI to cut any plasmids that lacked an activation domain variant and barcode insert, reducing the number of transformants with empty TFs without activation domains. We directed integrations to the *URA3* locus with a ~500 bp upstream homology region spanning the *URA3* promoter and *ACT1* promoter and a ~500 bp downstream homology region spanning the *TEF* terminator and *URA3* terminator. We plated the library transformations on YPD, grew overnight and replica-plated onto freshly prepared SC 5-FOA G418 plates (Amberg et al., 2005). We mated transformants en masse to an FY5 strain carrying the reporter integrated at the dubious ORF, YBR032w, and selected for diploids using on YfP with G418 (200 ug/ml) and NAT (100 ug/ml). The resulting diploids are prototrophs. The 20,000 yeast transformants were mated in batches. Before the final experiment, we pooled the batches and froze multiple aliquots. Before each sorting experiment, a frozen glycerol stock was thawed and grown overnight in SC+G418+NAT.

Strains were grown in synthetic complete (SC) dextrose media at 30°C (Amberg et al., 2005). Overnight cultures were diluted (1:5) into SC+1 uM ß-estradiol and grown for 3.5-4 hours at 30°C before FACS. For the amino acid starvation experiments, we washed the overnight culture with water twice and resuspened in minimal media without amino acids (synthetic dextrose, SD) with 1 uM B-estradiol for 4 hours before FACS.

### Fluorescence Activated Cell Sorting

We sorted the yeast library on a BD Aria-II machine. We used a yeast strain with the reporter and a TF lacking an activation domain as a negative control to determine autofluorescence and baseline mCherry levels. We sorted the library TF mutant containing cells into 8 bins: bin 1 had autofluorescent GFP levels (less than 95th percentile) and low mCherry (lower than 5th percentile); bin 2: autofluorescent GFP levels and normal mCherry levels; bins 3-8 all had greater than autofluorescent GFP levels, and evenly partitioned the distribution of GFP/mCherry ratios. We sorted >1 million cells into each bin. Sorted cells were grown overnight in SC, before gDNA was extracted (Zymo YeaSTAR D2002) and barcodes amplified by PCR (F: TGATAACTAGCTGA**GGGCCC**G R: GCTAT**GAATTC**GGACGAGGCAAGCTAAACAG NEB Q5 for 20 cycles) for sequencing on the Illumina NextSeq platform. We added PE2 indexing barcodes and phased P1 barcodes by digestion of barcode amplicons with ApaI and EcoRI and ligation of annealed adaptors. The final enrichment PCR used 4 ng of template and 20 cycles (NEB Phusion).

We analyzed individual constructs on a Beckman Coulter Cytomics FC 500 MPL with GFP only and processed the data with FlowJo 8 and Matlab.

### Sequencing analysis

After demultiplexing we kept all reads that perfectly matched our designed barcodes and flanking restriction sites. First, we normalized reads by the total counts in each bin and then, for each construct, renormalized abundance across bins. To convert the relative abundance across bins into an expression value, we computed a weighted average using the median GFP/mCherry ratio of each bin. Barcode abundance in the unsorted population had no correlation with computed activity (Fig S1D). The dynamic range is set by the GFP/mCherry ratios of the lowest and highest bins. For example, all constructs found only in the highest bin have the same weighted average activity. In the figures, we present the log_2_(activity/ WTmedian) in complete media (SC). The induction ratio in Figure 3 is the activity under starvation (SD) divided by the activity in SC (i.e. fold change during starvation). The unnormalized change in activity plotted in Figure S3B is the activity in SD minus the activity in SC.

We filtered out barcodes that we suspected to be attached to incorrectly synthesized activation domain variants. Based on the sanger sequencing of 91 colonies, we found two misassigned barcodes (template switching) and 55 perfect constructs (60%). To enrich for barcodes attached to the correctly synthesized activation domain variants, we amplified the full activation domain plus barcode region, performed paired-end sequencing of the unsorted yeast pool (2x150 bp, 2.1 M reads), merged reads with FLASH (Magoč and Salzberg, 2011) and looked for perfect constructs. For this sequencing library, we amplified the activation domain and barcode region with primers atatGGATCCctgcgggctctacttcatc and cgcggaattcTCGCTTATTTAGAAGTGGCG and digested with BamHI-HF and EcorI-HF. We created two filtered sets. Our high stringency set contained all barcodes for which the perfect construct was detected in the paired-end sequencing. By this analysis, we detected the designed construct for ~75% of barcodes. We used this set for all the analysis presented in the paper. In principle, this analysis could underestimate errors if a barcode is attached to two different activation domains. We also identified a very high stringency set where the majority of barcodes in the paired-end reads in the unsorted population were attached to the correct variant. We determined this second analysis overestimated errors because of template switching during the construction of the paired-end sequencing library. For the control sequences, a comparison of the full data, high stringency and very-high stringency data sets is shown in Figure S2A.

### Reproducibility analysis

We assessed reproducibility using the full library. We thawed glycerol stock aliquots, grew and sorted cells on separate days. The Pearson correlation between the two replicates was 0.988 and the Spearman rank correlation was 0.980. In replicate one, we sequenced 8-12 million (M) reads per bin and detected over 5,000 of the 6,500 designed barcodes. 3,888 barcodes had >100 reads across the 8 bins, our empirically determined cutoff for reliable measurement. In replicate 2, we sequences 18-23 M reads per bin and detected 5,930 constructs, 3,938 of which had >100 reads. For all analyses of activity in complete media, we used data from Replicate 1 only.

In subsequent experiments, we performed less sequencing. To determine how many reads were necessary, we resequenced Replicate 2 to a depth of 816 K reads. With a detection threshold of 10 reads, we measured 3,430 (87%) of the barcodes and found that the Pearson R = 0.989 and Spearman R = 0.981 compared to original Replicate 1. From this analysis we determined that much less sequencing was necessary for subsequent experiments. For the amino acid starvation experiment, we sequenced 2.4 M reads and used a detection cutoff of 10 reads. For the mCherry-only sort, we sequenced 5.9 M reads. In a typical run >87% of raw reads passed demultiplexing and size selection, and >93% of these could be matched to barcodes (>84% of raw reads). For a subset of the library, we compared overnight growth of sorted cells with direct extraction of DNA from sorted cells and found both methods gave very similar results (Pearson = 0.992; Spearman = 0.992; Fig S1B) after sequencing 1.2 M and 1.9 M reads respectively.

## QUANTIFICATION AND STATISTICAL ANALYSIS

### Data Analysis

Peptide sequence analysis was performed with custom python scripts using localCIDER (Holehouse et al., 2017) and PANDAS (McKinney and Others, 2010). All custom code was recorded in Jupyter Notebooks and is available upon request. Plots were created with matplotlib and Adobe Illustrator. Colors adapted from colorbrewer.org. Sequence frequency logos were created with WebLogo (Crooks et al., 2004).

To calculate the resolution of our measurement, we used the control sequences with multiple barcode sequences. For all possible pairs, we performed a two-sided, two sample t-test and report the smallest difference that was statistically significant at p < 0.05 after Bonferroni correction.

In Figure S2B, we searched for K-mer sequences that were enriched or depleted in the top or bottom 200 variants with a one tailed Fisher’s exact test (hypergeometric), using the frequency of the Kmer in the measured variants as the null. We removed orthologs and duplicate control sequences. We used a p-value cutoff of .01 with a Bonferroni correction for 44 positions and 200 sequences (p<.01/ (200*44)). Since the vast majority of our mutant sequences keep the WxxLF motif intact, we are statistically underpowered to detect enrichment of the primary motif in the top 200 sequences.

For the ANOVA analysis in Table S1, we used the *ols* module from *statsmodels.formula.api* in python. We used the Disorder and HoldChargeDisorder (D>E and E>D substitutions) variants. We report the adjusted R^2^. The models were: ‘NormExpS11 ~ Disorder,’ ‘NormExpS11 ~ Ws + Ys + Ls + Fs + Ms.’ and ‘NormExpS11 ~ Ws + Ys + Ls + Fs + Ms+Disorder.’

To determine if the parameters measured in the simulations were significantly different between the high and low activity sets, we performed permutation tests. We shuffled the sequence labels, keeping together the blocks of simulations for each sequence. For each permutation, we split the data into two groups and calculated the difference in means. Under this scheme, the differences in helicity were not significant. In 100,000 permutations, we never observed a difference in gyration radius or in WFY accessibility larger that than observed for the real data, supporting our assertion that these differences are statistically significant. Conversely, for helicity around the primary motif, p = 0.13, indicating the differences were not statistically significant.

In Figure 6A, to determine if adding a W in the secondary motif decreased activity was statistically significance, we shuffled the data. We shuffled the positions 100,000 times and calculated the mean activity of five consecutive positions to determine the empirical distribution mean values. In 100,000 trials, 658 random sets had a lower mean activity than the actual positions in Motif 2 [positions 7,8,9,10,11,12] leading us to an empirical p = 0.00658. Similarly, 67 random sets of five have a greater mean activity than the actual value of the positions around Motif 1 [positions 21,22,25,26,27], leading to an empirical p = 0.00067.

In Figure 6D, to determine which positions were under-mutated or over-mutated in the most induced variants, we first used Fisher’s exact test. We looked at the 100 mutants with the largest raw increase in activity. We created a matched set of low induction mutants with similar range of activity levels: we selected the 100 least inducible mutants that have activity within the same range as the most induced mutants, to separate mutations that lead to high activity from mutations that lead to moderate activity and low induction. No individual position was mutated more than expected, (p <0.05, Bonferroni corrected for 44 positions). To test if the set of six central acidic residues were under-mutated, we used a permutation test. Using the control, charge, disorder and hydrophobicity mutants, we sampled sets of 100 sequences and computed the average number of times those six positions were mutated. In 10^5^ permutations, only one instance had a lower average mutation load than the actual average number.

### Machine Learning

For the support vector regression, we used the sklearn python package (Pedregosa et al., 2011). We included all the control, disorder, hydrophobicity and charge variants in the high confidence set (N= 1738). We normalized the magnitude of all input parameters to [0,1]. For the SVR we used the gaussian radial basis function as the kernel, and set C=1e3 and gamma=0.1. To expedite model exploration, we fit and tested on all the data. We reexamined high performing models with five-fold cross validation and report the average R^2^ of multiple cross validation runs. Our initial model search explored all combinations of counting one to five amino acids. The second round search retained amino acids frequently found the highest scoring models and adding net charge calculated over the whole peptides or subregions. Once we determined that W, F, Y, M and L were consistently in the top models, we fixed these parameters and tried adding the local net charge calculated over each possible window of 5-44 amino acids at all positions. We ultimately chose to calculate net charge over positions 12-25, but this window is only marginally better than similar windows.

To apply this model to all 44 amino acid regions of proteins with annotated DBDs, we downloaded a list of 162 proteins from YeastMine (yeastmine.yeastgenome.org) and manually added ZAP1. We chopped this list to 44 amino acid regions and computed the 6 parameters of each window. We normalized this matrix by the same scaling factors used to normalize the Gcn4 training data. We applied the model to each window. 49% of windows scored more highly than WT Gcn4.

To sample random peptides from the yeast genome, we used the non-redundant list of S. cerevisiae reviewed proteins taken from UniProt (UP000002311_559292). We concatenated all these proteins and sampled 50K nonoverlapping regions of length 44. Specifically, we picked a pseudorandom integer from [0-10], took the next 44 residues, then picked another randoms spacer.

### Simulation methods

All simulations were run with the CAMPARI Monte Carlo simulation engine (http://campari.sourceforge.net) and the ABSINTH implicit solvent paradigm (Vitalis and Pappu, 2009). The combination of CAMPARI and ABSINTH has been used to characterize the conformational ensemble of a wide variety of IDRs, providing quantitative agreement with a variety of experimental measures (Martin et al., 2016; Metskas and Rhoades, 2015; Vitalis and Pappu, 2009). We simulated the 50 most active and least active variants from the charge, hydrophobicity and disorder mutation sets (300 unique sequences). In addition, we simulated the 50 most induced variants (by unnormalized change in activity) and a set of variants that were not induced. The uninduced set was selected to span the same range of activities as the 50 most induced variants, which we used to separate ‘low induction’ signals from ‘high activity’ signals. Simulation snapshots were visualized and overlaid with VMD (Humphrey et al., 1996).

Each simulation was started from a random, non-overlapping conformation and run for 50 M steps (40 M production), with ten independent simulations per sequence (18 for WT), giving a total of 5,008 independent simulations. Conformations were extracted every 5,000 steps, generating 8,000 conformations per simulation (80,000 per sequence). Simulations were analyzed using MDTraj (McGibbon et al., 2015) and CTraj (http://pappulab.wustl.edu/CTraj.html). Y/F/W accessibility is determined by calculating the summed solvent accessible surface area of Y, F and W residues with a 1.5 nm spherical probe, which we treat as a proxy for a protein interaction surface. Varying the probe-size to 1.0 nm rescaled the absolute values but did not change the results. Helicity was calculated using the DSSP algorithm (Kabsch and Sander, 1983). The radius of gyration is calculated as the instantaneous mass-unweighted distribution of atoms in each conformation, as used previously (Holehouse et al., 2015).

To determine if the addition of arbitrarily placed W residues will inevitably lead to an increase in the accessibility of Y/F/W residues we ran simulations of a Gcn4 length-matched (44 AAs) sub-region of a different intrinsically disordered region Ash1. The Ash1 intrinsically disordered region has been extensively studied by small angle X-ray scattering, nuclear magnetic resonance and simulations; it is known to be disordered with a strong propensity to be highly expanded (Martin et al., 2016). This protein is an ideal test-case; if the W residues have no impact on the conformational behaviour of the intrinsically disordered region then we would expect a large increase in the Y/F/W accessibility in response to additional W residues. We randomly placed one, two, or three W residues in the backdrop of the central region of Ash1 (sequence: MRSRSSSPVRPKAYTPSPRSPNYHRFALDSPPQSPRRSSNSSI), randomly selecting three different positions such that there are three distinct sequences with one, two, or three additional W residues. We then ran simulations in triplicate and determined the Y/F/W accessibility (Fig S4D). Error bars describe the standard error of the mean. We found that the number of W residues added was a poor determinant of the Y/F/W accessibility; the number of W residues added to Ash1 had no correlation with the Y/F/W accessibility.

### Sequence analysis

Proteome-wide disorder analysis was performed by identifying long (>20 residue) disordered regions using the IUPred algorithm with a threshold of 0.4 (Dosztanyi et al., 2005). This analysis yields results consistent with previously performed proteome-wide analysis using meta-predictions from the D2P2 web-server (Holehouse et al., 2017; Oates et al., 2013). Analysis of disordered regions was performed using localCIDER (Holehouse et al., 2017).

## Acknowledgements

We thank Gary Stormo, Ashley Wolf and members of the Cohen Lab for feedback on the manuscript and helpful discussions. We thank Jessica Hoisington-Lopez for support with high-throughput sequencing. M.V.S. holds a Postdoctoral Fellowship from the American Cancer Society and a Postdoctoral Enrichment Program Award from the Burroughs Wellcome Fund. This work was supported in part by the Washington University Bonnie and Kent Lattig graduate fellowship (A.S.H.), National Institutes of Health (NIH) grants R01-GM092910 and R01-HG008687 to B.A.C., NIH grant 5RO1-NS056114 and National Science Foundation grant MCB1614766 to R.V.P.

## Author Contributions

M.V.S. and B.A.C. conceived the project. M.V.S., B.A.C, R.D. and R.V.P. designed the mutations. D.S. created part of the yeast library. M.V.S. performed experiments and analyzed the data. A.S.H. ran and analyzed the simulations. M.V.S. and B.A.C. wrote the paper with input from all the authors.

## DATA AND SOFTWARE AVAILABILITY

We have included the following supplemental data tables

### Data S1: Activation Domain Activity and physical properties

Columns are: Index, ActivationDomainSeq (protein sequence of designed variant), ActivityCompleteMediaReplicate1 (Raw activity), ActivityCompleteMediaReplicate2 (Raw activity), ActivityCompleteAAStarvation (Raw activity), MutationClass, HighStringencySet (Boolean), VeryHighStringencySet

Index, ActivityCompleteMediaReplicate1_Normalized, ActivityCompleteMediaReplicate2_Normalized, Induction (ratio of ActivityAAStarvation_Raw to ActivityCompleteMediaReplicate1_Raw), ActivityCompleteMediaReplicate1_Raw, ActivityCompleteMediaReplicate2_Raw, ActivityAAStarvation_Raw, MutationClass,Charge, Hydrophobicity (normalized Kyte Doolittle scale), Kappa (Das and Pappu, 2013), Disorder (IUpred prediction), HighStringencySet (Boolean), VeryHighStringencySet (Boolean), mCherryOnly_Raw

*MutationClass* is the category assigned to each mutant: Control, Charge, Hydrophobicity, Kappa (Shuffle mutants, initially designed to systematically vary the Kappa parameter), MovePrimaryMotif, DirectOrthologs, ChopOrthologs, Disorder, HoldChargeDisorder (D>E and E>D substitutions).

### Data S2: Barcode Counts

This table contains the raw barcode counts for each activation domain in each bin of each experiment. There are 4 experiments each with 8 bins: S11 is replicate 1 in SC, S12 is replicate 2 in SC, S14 is amino acid starvation, and S15 is grown in SC and sorted on mCherry only. Note that the GFP/mCherry Ratio of each bin is included in the column header. For the mCherry bins in S15, the value of mCherry is included in the column header.

### Data S3: Orthologs

A manually collected set of sequences orthologous to Gcn4 used for the library construction.

### Data S4: 44 amino acid regions of the yeast proteome with high predicted intrinsic disorder, high acidity and many hydrophobic residues

We computationally chopped the non-redundant yeast proteome into all 44 amino acid windows. We selected for windows with IUpred >0.4, more net acidity than Gcn4 and more hydrophobic residues (W,F,Y,L,M,I,V) than Gcn4. IUpredlong was computed on the full protein sequences and averaged over each window.

